# Circling in on plasmids: benchmarking plasmid detection and reconstruction tools for short-read data from diverse species

**DOI:** 10.1101/2025.07.28.667252

**Authors:** Marco Teixeira, Celia Souque, Colin J. Worby, Terrance Shea, Nicoletta Commins, Joshua T. Smith, Arjun M. Miklos, Thomas Abeel, Ashlee M. Earl, Abigail L. Manson

## Abstract

The ability to detect and reconstruct plasmids from genome assemblies is crucial for studying the evolution and spread of antimicrobial resistance and virulence in bacteria. Though long-read sequencing technologies have made reconstructing plasmids easier, most (97%) of the bacterial genome assemblies in the public domain are generated from short-read data. Work to compare plasmid reconstruction tools has focused primarily on *E. coli*, leaving gaps in our understanding of how well these tools perform on other, less well-characterized, taxa.

Using high quality assemblies as ground truth, we benchmarked 12 plasmid detection tools (which identify plasmid contigs in assemblies) and four plasmid reconstruction tools (which group contigs from the same plasmid together). We tested their ability to characterize diverse plasmids from short-read assemblies representing a wide range of Enterobacterales and *Enterococcus* species, including newly discovered and poorly characterized species collected from non-human hosts. Plasmer, PlasmidEC, PlaScope, and gplas2 were the highest-scoring plasmid detection tools, performing well for both Enterobacterales and enterococci. The two major determinants of accurate plasmid detection were representation in plasmid databases - with Enterobacterales plasmids being more easily detected than those from enterococci - and assembly contiguity, which was also key for successful plasmid reconstruction. Gplas2 performed best for plasmid reconstruction; however, less than half of plasmids were perfectly reconstructed, suggesting that substantial room for improvement remains in this class of tools.

**Key Messages:**

- The ability to detect and reconstruct plasmids from short-read assemblies is crucial to study the spread of antimicrobial resistance and virulence genes in bacteria.
- Most past comparisons of tools for plasmid detection and reconstruction have focused on plasmids from well-characterized *E. coli*; therefore, we broadened our benchmarking set to include plasmids from diverse Enterobacterales and *Enterococcus* species, collected from a wide range of hosts.
- We compared the predictions of 12 recent plasmid detection and four plasmid reconstruction tools on short read assemblies, against a truth set generated from high-quality hybrid assemblies.
- Plasmer, PlasmidEC, PlaScope, and gplas2 were the highest-scoring plasmid detection tools.
- Gplas2 was the highest-scoring tool for plasmid reconstruction; the quality of plasmid reconstruction was mainly determined by assembly fragmentation, with plasmids in more contiguous assemblies being easier to reconstruct.

## Introduction

Plasmids play a major role in the evolution and spread of antimicrobial resistance and virulence in bacteria, as they frequently contain antimicrobial resistance genes (ARGs) and factors that contribute to bacterial pathogenicity [1, 2, 3, 4, 5, 6]. However, deciphering plasmid content in a bacterial genome assembly can be challenging, particularly for assemblies generated from short read datasets, which represent 97% of publicly available data [7]. Repeat regions and insertion sequences can be shared with other plasmids or the chromosome, and frequently lead to breaks in short read assemblies [8]. Recombination between plasmids leads to rapidly evolving mosaic structures that are highly variable between organisms, meaning that newly-sequenced plasmids may not be represented in gene content and synteny within existing plasmid databases [9]. Even long-read sequencing technologies, which are known for creating datasets that are more amenable for plasmid reconstruction, can have blind spots, missing small plasmids [10].

Current tools for characterizing plasmids from short-read assemblies can be broadly divided into those that detect plasmids and those that reconstruct plasmids. Generally, plasmid detection tools predict whether a contig represents part of a plasmid or a chromosome, while plasmid reconstruction tools (also referred to as plasmid contig binning tools) group together contigs from the same plasmid or chromosome.

Past comparisons have focused on plasmid reconstruction tools, mainly for *E. coli* [11, 12], an extensively studied species that has contributed the most to plasmid reference databases used by many plasmid detection tools [13]. One such large, recent benchmark of six tools [12] found that the majority of *E. coli* plasmids carrying ARGs could not be completely reconstructed by any tool, and that large ARG-containing plasmids were the most difficult to reconstruct. While MOB-suite was the highest-performing overall plasmid reconstruction tool, plasmidSPAdes performed best for small plasmids, hinting that the best tool depends on the characteristics of the plasmids being reconstructed. MOB-suite also outperformed other plasmid reconstruction tools in a separate, comparative, *E. coli* -focused study, though the authors noted the need for improved plasmid reconstruction tools [11].

Motivated by interests in evaluating the performance of plasmid detection on organisms other than *E. coli*, and assessing recently developed plasmid reconstruction tools not considered by previous benchmarking efforts, we undertook a systematic benchmarking of 13 tools (of which 12 are suitable for plasmid detection and four for plasmid reconstruction) on a diverse set of 250 genomes from Enterobacterales and *Enterococcus* species.

Studying the performance of these tools across different genera, including one that has contributed relatively little to popular plasmid databases, as well as their strengths and pitfalls, will improve the community’s ability to leverage the millions of short-read datasets currently available [7] and provide a better understanding of the mechanisms of spread of antibiotic resistance.

## Methods

### Generating a benchmarking dataset from diverse isolates

We generated Oxford Nanopore (ONT) long-read sequencing data for 93 Enterobacterales hospital isolates [14, 15], and 157 *Enterococcus* isolates collected from diverse animals and remote environments [16, 17], some presented here for the first time (Supplementary Table 1). All isolates were also sequenced using the short-read Illumina platform (Supplementary Table 1).

**Table 1:**
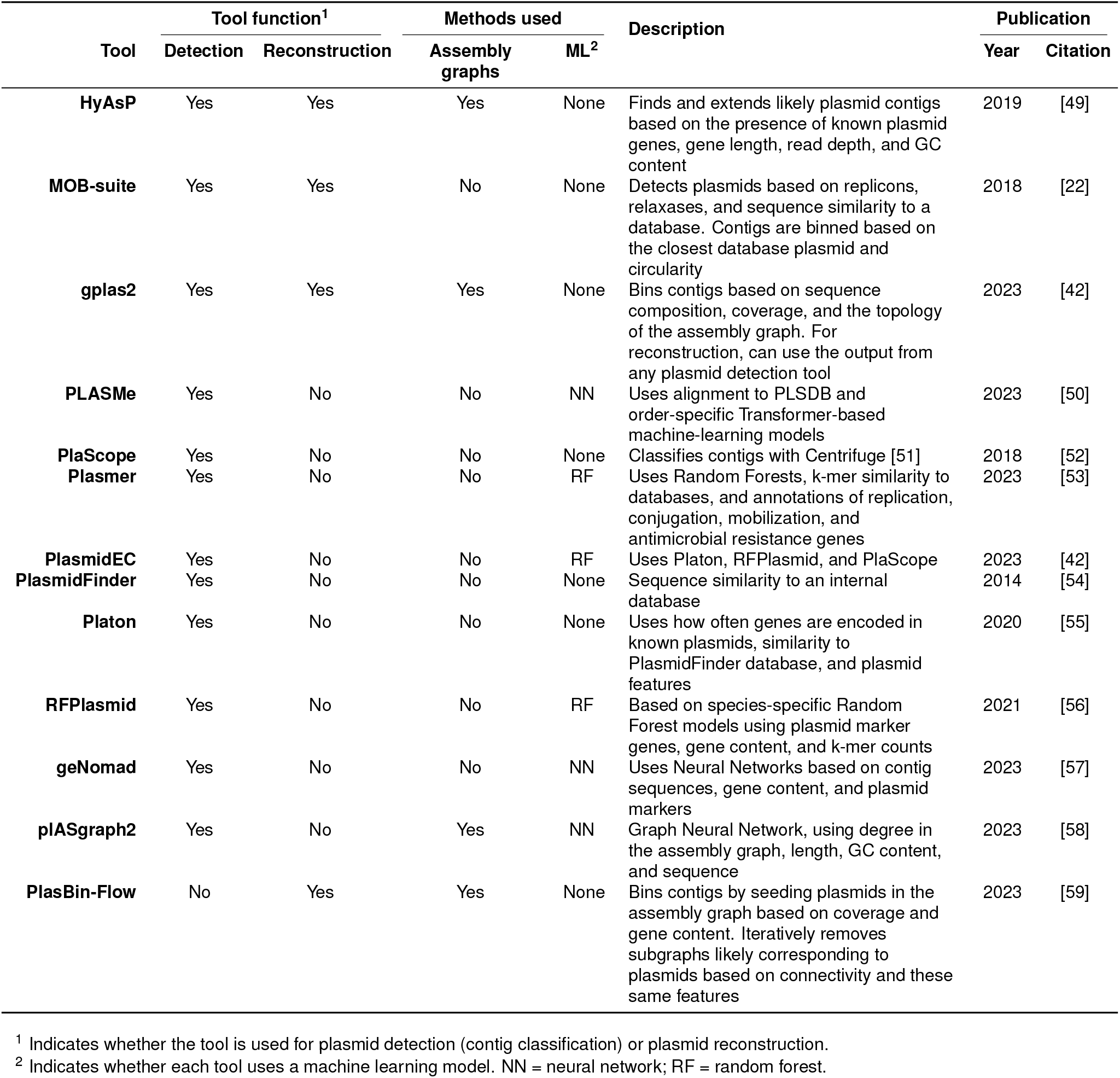
Tools tested in benchmarking.

High-quality hybrid assemblies were generated for each isolate by combining both short and long reads. Hybrid assemblies were generated both with all sequence data, and with ONT data subsampled to 50x, following methods previously described [18, 15], using Unicycler v0.4.4 (with default settings) and refinement with Pilon v1.23 [19]. Out of the two hybrid assemblies, the more contiguous one was selected to determine ground-truth contig annotations. GAEMR (https://github.com/broadinstitute/GAEMR) was used to align contigs to the NCBI Nucleotide database using discontiguous Mega BLAST [20]; the resulting metrics were used to remove 276 contaminant contigs in 36 of the enterococcal hybrid assemblies.

Short-read-only assemblies were also generated with Unicycler v0.5.4 [21], using default settings, which were used as inputs for the plasmid detection and reconstruction tools evaluated. We used Unicycler as it generates the assembly graph required for one of the tools evaluated (gplas2). Contigs with fewer than 1000 bp were excluded when evaluating tools.

### Establishing ground-truth plasmid annotations

The high-quality hybrid assemblies were used to establish ground-truth plasmid annotations. For hybrid assemblies in which all contigs were labeled as circular by Unicycler, we defined the largest contig as the chromosome and all other contigs as plasmids. When not all contigs were labelled as circular, we inspected each contig manually to determine the correct plasmid annotation, including looking at its BLAST hit against the NCBI Nucleotide database and the relative plasmid copy number indicated by read coverage.

In order to annotate the replicon cluster for each plasmid in complete hybrid assemblies, we used the MOB-typer module of MOB-suite v1.4.9 (database v3.1.8) [22]. Enterobacterales plasmids were further grouped into incompatibility groups, according to their replicon cluster annotations from MOB-suite, as these groups are more easily interpretable and commonly used in the literature. As few enterococcal plasmids had replicons from characterized incompatibility groups, the original replicon cluster annotations were used for enterococci. Plasmids assigned to multiple replicon clusters were characterized as multi-replicon, whereas those with uncharacterized replicon clusters were characterized as “unknown incompatibility group”.

The ground-truth plasmid annotation for each short-read contig (chromosomal or plasmid) was determined by aligning them to their corresponding high-quality hybrid assemblies using nucmer (MUMmer v3.23, [23]). Each short-read contig was labeled as plasmid or chromosomal based on the best alignment to the hybrid assembly; contigs mapping equally well to a hybrid-assembly chromosome and plasmid contig were excluded from downstream analyses. For complete hybrid assemblies (with all contigs labelled as circular), short-read contigs aligning to the same hybrid contig were presumed to be from the same plasmid/chromosome. Incomplete hybrid assemblies were excluded when comparing plasmid reconstruction tools, as it is not possible to determine a ground-truth.

### Assessing plasmid and chromosomal sequence diversity in our isolate collection

To quantify the chromosomal sequence diversity of the isolates in our collection, we calculated all-vs-all Average Nucleotide Identity (ANI) between the chromosomal content of our complete hybrid assemblies, computed with fastANI [24] version 1.32, using a k-mer size of 16, a minimum alignment fraction of 0.2, and a fragment length of 3000.

Similarly, in order to quantify the plasmid sequence diversity within our collection of isolates with complete hybrid assemblies, we computed the ANI between each plasmid, with (greater than the 0.2 used for chromosomes) as mobile genetic elements often lead to high ANI between unrelated plasmids with low alignment fractions [25]. We also assessed plasmid diversity by clustering plasmids from complete assemblies according to their sequence similarity by performing Louvain clustering [26] based on the alignment fraction determined when calculating ANI. We selected the resolution parameter by testing values between 0.1 and 3 in increments of 0.1 and choosing the one that maximized modularity.

To assess plasmid database representation for our dataset, especially between taxa, we searched the plasmids in our complete hybrid assemblies against the PLSDB database [13] – one of the most comprehensive and popular plasmid databases – using Mash v2.0 [27] (sketch size of 1000). Plasmids were considered present in PLSDB if they had any hits with identity greater than 99.9% (over their full length). Furthermore, we assessed the identities between each plasmid and its best match in PLSDB according to the number of matching hashes.

### Evaluating the performance of plasmid detection and reconstruction tools

The short-read assemblies were used as input for the plasmid detection and reconstruction tools (Table 1). Details for how each tool was run can be found in Supplementary Table 2. Predictions for each short-read contig were compared to the ground truth plasmid annotations.

**Table 2:** Short-read assembly quality metrics.

|  | Median number of contigs (IQR)* |  |
| --- | --- | --- |
|  | Enterobacterales | Enterococcus |
| **Number of contigs** | 72 (58) | 26 (15) |
| **Chromosomal** | 53 (45) | 21 (14) |
| **Plasmid** | 21 (18) | 4 (6) |
| **N50** | 212 041 (176 561) | 303 098 (163 414) |
| **Assembly length** | 5 175 001 (414 196) | 2 965 356 (390 262) |

\* excluding contigs < 1kbp

Plasmid detection was evaluated by calculating precision, recall, and F1 score, defining true positives as plasmid contigs correctly classified, false positives as chromosomal contigs misclassified as plasmidic, and false negatives as plasmid contigs misclassified as chromosomal. Plasmid reconstruction was evaluated by calculating the Normalized Mutual Information (NMI) [28], homogeneity [29], and completeness [29]. These metrics have been widely used to evaluate clustering results when the ground-truth clustering assignments are known, a problem similar to the grouping of short-read contigs from the same plasmid or chromosome [30]. NMI is based on the entropy (*H*, Equation 1) of the set of ground-truth plasmids of origin (*S* = *{s*_1_, *s*_2_, …, *s*_*N*_*}*) and the bins assigned to each contig (*B* = *{b*_1_, *b*_2_, …, *b*_*K*_*}*), according to Equation 2, where *P* (*u*_*i*_) represents the probability of a contig being assigned to the *i*-th element of set *U*.

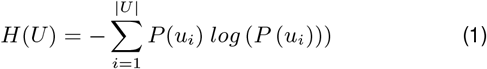

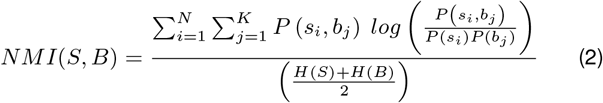

This metric takes a value of 100% when all contigs are grouped according to the plasmid of origin, while an NMI closer to 0% is indicative of poor reconstruction. Homogeneity (Equation 3) reflects the purity of the reconstructed plasmids - bins containing contigs from just one plasmid lead to a homogeneity of 100%, decreasing as contigs from different plasmids are placed together in the same bin.

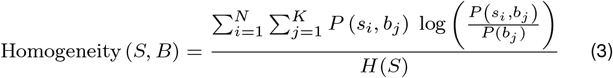

Completeness (Equation 4) quantifies how often plasmids were fragmented into different bins - placing all contigs from a plasmid in the same bin results in 100% completeness, and this measure takes lower values as more contigs from the same plasmid are placed in different bins.

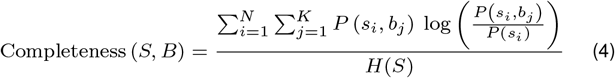

In some instances, short-read contigs may be assigned to multiple ground-truth plasmids or to multiple bins by the plasmid reconstruction tools. As NMI, homogeneity, and completeness cannot be defined in these cases, we took the cross-product of ground-truth plasmids and predicted bins before computing these metrics. For instance, a contig originating from plasmids *s*_1_ and *s*_2_ and assigned to bins *b*_1_ and *b*_2_ would be split into four similar contigs with (*s*_1_, *b*_1_), (*s*_1_, *b*_2_), (*s*_2_, *b*_1_), and (*s*_2_, *b*_2_) as their ground-truth plasmid and predicted bin, respectively.

For each of these metrics, we calculated the 95% confidence intervals through bootstrapping with 1000 iterations, sampling isolates with replacement. The 2.5 and 97.5 percentiles of the resulting metrics were taken as the limits of the 95% confidence interval.

### Assessing the effect of contig length on plasmid detection

We binned contigs from short-read Enterobacterales and enterococci assemblies according to their length. To ensure a similar number of contigs per bin, we binned them into three bins per log-scale division (i.e. 1-4 kbp, >4-7 kbp, >7-10 kbp, etc.). Then, we calculated the plasmid detection metrics separately for contigs in each bin. The correlation between the performance metrics and the length of the input short-read contigs was assessed by comparing the upper limit of the bin and the mean value of each metric after bootstrapping according to the Spearman correlation coefficient. The *p*-values refer to a two-sided permutation test (10 000 iterations) of the null hypothesis that the contig length and the plasmid detection metrics are uncorrelated.

### Comparing the detection performance for short and long plasmids

We classified plasmids from isolates with finished hybrid assemblies according to their size in the ground-truth hybrid assemblies. Enterobacterales plasmids longer than 30 kbp and enterococcal plasmids longer than 28 kbp were classified as “large”, consistent with previous classifications [31]; all others were classified as “small”. We then computed the recall for both groups as previously described.

### Determining the effect of antimicrobial resistance genes (ARGs) on plasmid detection and reconstruction

We annotated ARGs in both the short-read and hybrid assemblies with RGI v5.1.1, using the CARD v3.2.9 database [32], excluding genes associated with antibiotic efflux (as these are common and usually depend on other resistance factors to confer resistance [33]) and hits labelled by RGI as “loose”. We divided all short-read contigs into sets with and without at least one ARG, and compared the performance of plasmid detection tools between the two sets, using the performance metrics for plasmid detection as previously described. We also compared the quality of plasmid reconstruction for plasmids with ARGs versus for plasmids not encoding resistance. For this, we excluded all chromosomal short-read contigs and divided plasmids and their short-read contigs into ARG-containing and non-ARG-containing (depending on the ARG content of the corresponding hybrid contig). The plasmid reconstruction metrics were calculated for each group as described above.

### Disentangling the effects of specific assembly characteristics on plasmid detection and reconstruction

We used generalized linear models to disentangle the individual effects of specific assembly characteristics on plasmid detection and reconstruction. We only included isolates with a complete hybrid assembly so as to accurately determine plasmid characteristics such as length and presence in plasmid databases. Logistic regression and linear regression models (statsmodels package, v0.14.2) were trained on 80% of the short-read contigs and assemblies, respectively, with the remaining 20% used as a test set to determine generalizability. Short-read contigs were split into the training and test sets, stratifying by contig annotation (plasmid or chromosomal).

For each of the four best plasmid detection tools (PlaScope, PlasmidEC, Plasmer, and gplas2), we applied two logistic regression models (one trained with Enterobacterales assemblies and the other trained with enterococcal assemblies) to estimate if that tool correctly predicted plasmid annotations (chromosomal or plasmid) for each short-read contig. We examined contributions from the following covariates: i) the length of the short-read contig (in Mbp); ii) whether the short-read contig contained ARGs; iii) the number of transposases in the entire hybrid assembly; iv) plasmid size, relative to small plasmids (contigs were grouped into three classes: chromosomal, small plasmid, and large plasmid); and v) how similar the corresponding plasmid in the hybrid assembly was to its closest representative in the PLSDB database, quantified as the Mash identity between the corresponding hybrid contig and the PLSDB hit with the most matching Mash hashes. Chromosomal contigs (without matches to PLSDB) were assigned the median identity across all plasmid contigs, minimizing the correlation between this identity and contig annotations.

We also used a linear regression model to predict gplas2’s overall plasmid reconstruction quality, quantified with the NMI, for each short-read assembly based on the following covariates: i) the fragmentation of the short-read assembly, quantified as the average number of short-read contigs per hybrid contig (i.e., per plasmid and chromosome); ii) the total number of plasmids and chromosomes in the hybrid assembly; iii) the number of transposases in the hybrid assembly (i.e., the entire genome), counting multiple copies of the same transposase; and iv) the number of short-read contigs with ARGs. Samples for which gplas2 found no plasmids (n=25) were discarded when fitting the model.

We annotated transposases in the hybrid assemblies by aligning them to the TnCentral and ISFinder databases (downloaded on 2024/10/07) [34, 35] with BLASTn 2.2.30 [36]. For overlapping hits (including partial overlaps), we kept the alignment with the greatest identity.

## Results

### Our benchmarking dataset contained 475 diverse plasmids from 31 species from clinical and environmental settings

In order to evaluate the performance of plasmid detection and reconstruction tools, we first generated truth sets of high-quality hybrid assemblies (truth assemblies) from 93 clinical Enterobacterales, representing 14 species from six genera, and 157 mostly environmental *Enterococcus* spp., representing 17 species (Figure 1A, Supplementary Table 3) [14, 15, 16]. Many of the datasets used to create the truth assemblies (Supplementary Table 1) were new to this study and none of the assemblies were in the databases used by the tools evaluated, which could otherwise lead to overestimating their performance for novel plasmids.

**Figure 1.**
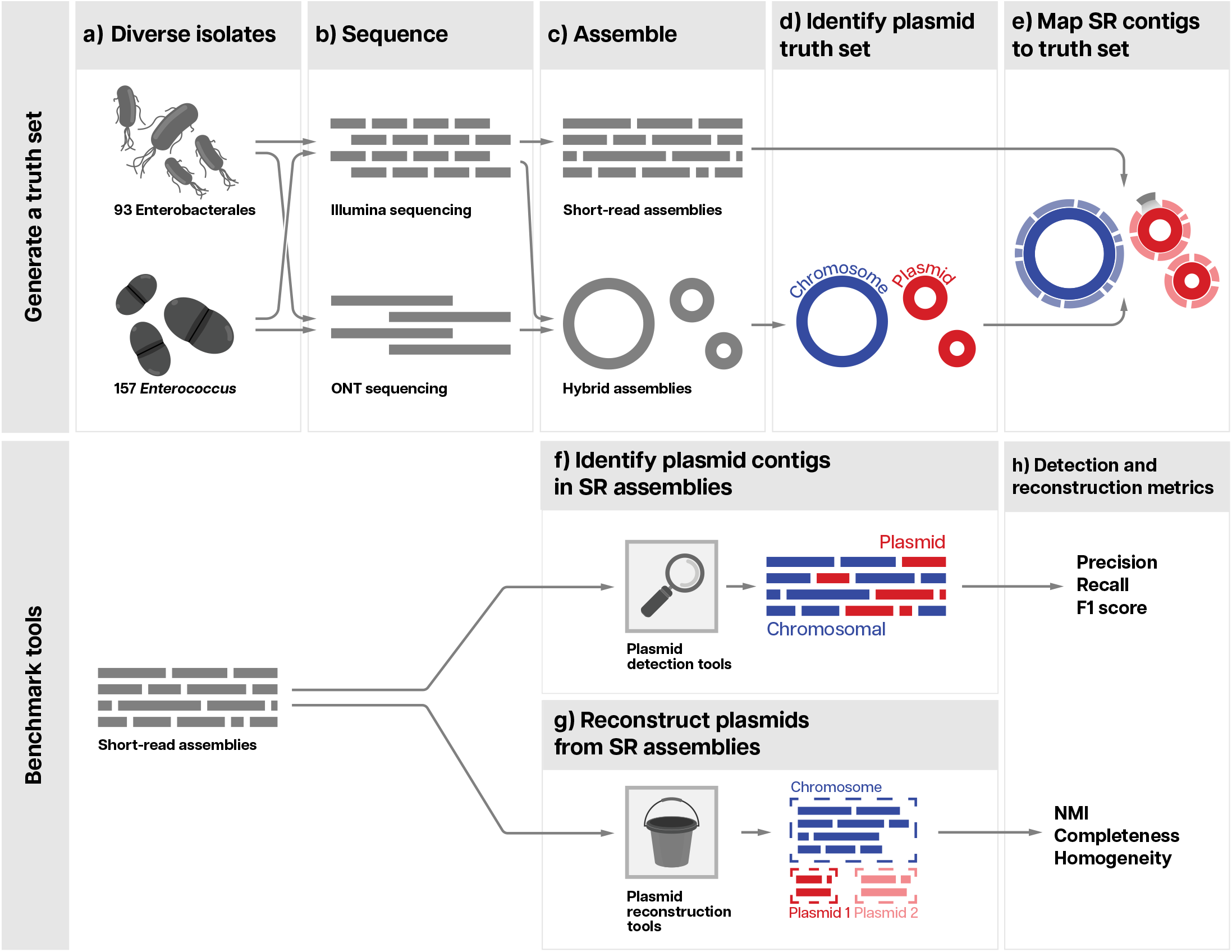
Benchmarking plasmid detection and reconstruction tools. **a)** We selected 93 Enterobacterales and 157 enterococcal isolates with diverse species and plasmids; **b)** sequenced them with short-read (SR) Illumina and long-read ONT technologies; and **c)** generated SR and high-quality hybrid assemblies with Unicycler. **d)** We classified contigs in the hybrid assemblies as chromosomal or plasmidic to establish a truth set (Methods), then **e)** mapped SR contigs to the truth set. We used SR assemblies as input for **f)** plasmid detection tools, which classify contigs as chromosomal or plasmidic, and **g)** plasmid reconstruction tools, which bin together contigs originating from the same plasmid or chromosome. **h)** We assessed the performance of tools based on the metrics indicated.

All 157 *Enterococcus* and 48 (52%) of the Enterobacterales truth assemblies consisted of only circular contigs, with a median of two and three plasmids per isolate for enterococci and Enterobacterales, respectively. The remaining 45 Enterobacterales assemblies had a median of 8 contigs, of which 45.8% were circular. In total, the truth assemblies represented 171 Enterobacterales and 304 *Enterococcus* plasmids with overall high diversity: 45 and 78 clusters based on gene content similarity from Enterobacterales and enterococci, respectively (Supplementary Table 4, Figure 2, Methods).

**Figure 2.**
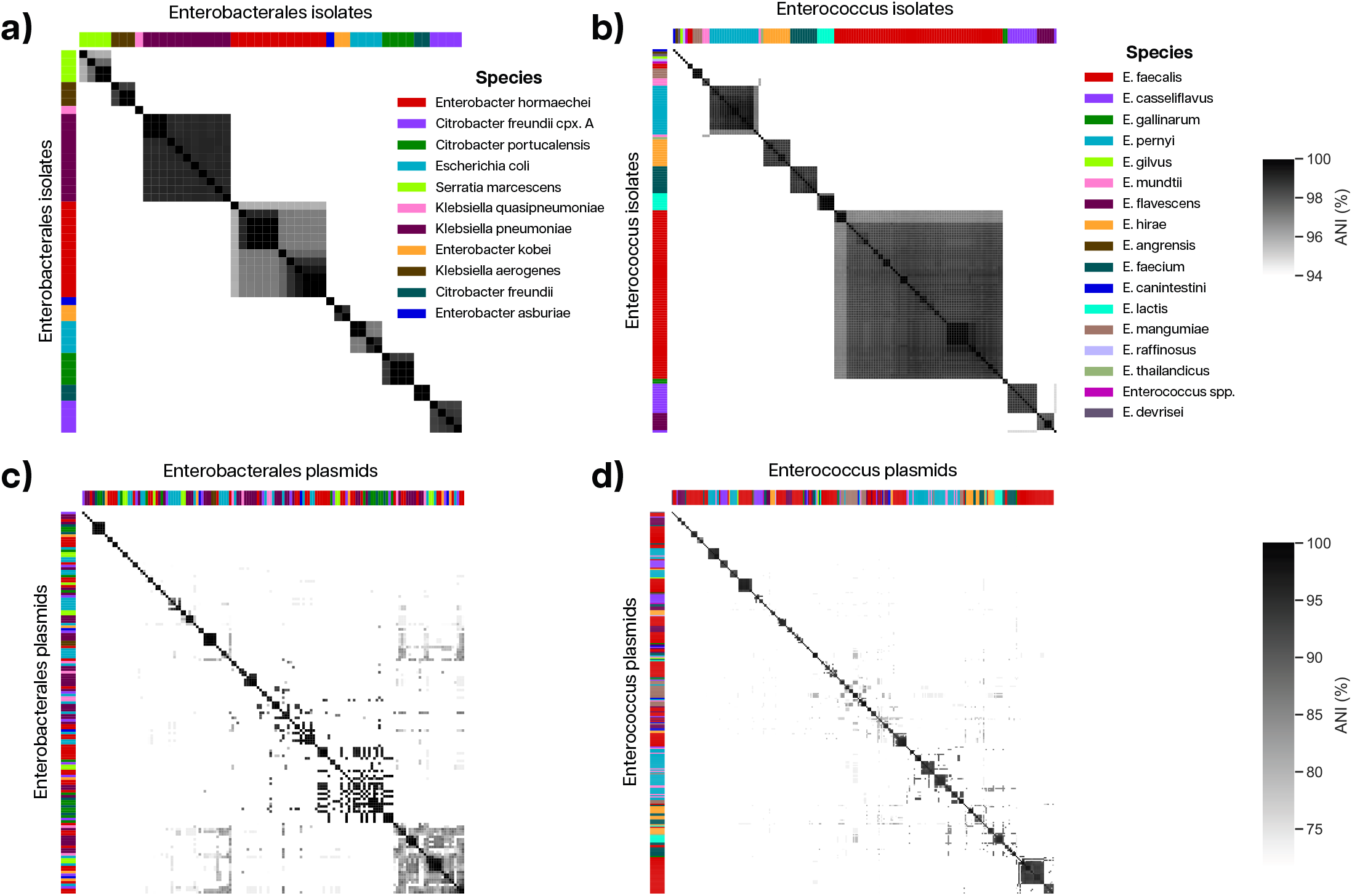
Our dataset included diverse isolates and plasmids. **a-b)** Pairwise ANI between same-species isolates from **a)** Enterobacterales and **b)** *Enterococcus*. ANI values > 94% are shown. **c-d)** Pairwise ANI between **c)** Enterobacterales and **d)** enterococcal plasmid pairs with an alignment fraction > 50% in finished assemblies. Strips on the axes are colored according to species of origin. ANI values > 75% are shown.

Of the *Enterococcus* and Enterobacterales plasmid sequences, 75% and 11%, respectively, were not found in the popular plasmid database PLSDB [13] (at 99.9% Mash identity, including partial matches), reflecting the bias in plasmid databases towards more well-studied and clinically related species. Overall, the average identity to the best PLSDB match was 85.3% for *Enterococcus* plasmids and 95.3% for Enterobacterales plasmids (Supplementary Figure 1).

We also generated short-read Illumina-only assemblies for each isolate in the truth assembly set. As expected, the short-read assemblies were more fragmented (Supplementary Table 1, Supplementary Table 3) than the hybrid ones. Overall, the *Enterococcus* short-read assemblies were more contiguous than those from Enterobacterales (N50 of 303 098 vs. 212 041), and had a smaller total assembly size (Table 2).

To determine whether a contig from the short-read assembly was part of a plasmid or the chromosome, we aligned each contig to its respective truth assembly. The short-read assemblies captured most of the plasmid content: a median of 96.1% (IQR=13.9%) and 97.9% (IQR=9.8%) of Enterobacterales and *Enterococcus* plasmid base pairs, respectively. The plasmids in short-read assemblies tended to be split into multiple contigs, with those from Enterobacterales being more fragmented than those from the enterococci (a median of 2 (IQR=7) and 1 (IQR=3) short-read contigs per plasmid, respectively). 49.7% (n=314) of *Enterococcus* and 30.4% (n=227) of Enterobacterales plasmids were fully captured in a circular contig.

### Comparing plasmid detection and reconstruction tools

We next compiled a set of 12 tools for detecting plasmids and 4 tools for reconstructing plasmids from short-read genome assemblies (Table 1), focusing on recently developed tools (published between 2018 and 2023), as well as the most highly cited plasmid detection tool: PlasmidFinder. An in-depth comparison of older tools was published by Paganini et al. [12]. We did not include tools for metagenomic analyses (e.g. SCAPP [37] 4 and metaplasmidSPAdes [8]). The tools we selected for benchmarking use different methods for detecting and reconstructing plasmids, including assembly graphs (HyAsP, gplas2, plASgraph2, and PlasBin-Flow), and machine learning (Plasmer, PlasmidEC, RFPlasmid, PLASMe, geNomad and plASgraph2) (Table 1). Each tool was run using the short-read assemblies as input (Supplementary Table 2).

#### Plasmid detection

##### PlaScope, PlasmidEC, Plasmer, and gplas2 were better at detecting plasmid contigs

Overall, different tools varied in their performance (Figure 3a, Supplementary Table 5). Most chromosomal (60.3%) but few plasmid (5.6%) contigs were correctly assigned by all tools. However, most contigs were correctly classified by at least one tool: just two chromosomal Enterobacterales and 20 plasmidic *Enterococcus* contigs were misclassified by all tools. Overall, the same four tools achieved the highest F1 metric across both Enterobacterales and enterococci: PlaScope, PlasmidEC, Plasmer, and gplas2. These tools correctly detected contigs representing on average 87%-99% of all base pairs in plasmids for Enterobacterales, and 80%-95% for *Enterococcus* (Supplementary Table 6). PlaScope performed slightly better for Enterobacterales, while Plasmer performed slightly better for the enterococci. The results from PlasmidEC - an ensemble of tools including PlaScope - were similar to those from PlaScope alone.

**Figure 3.**
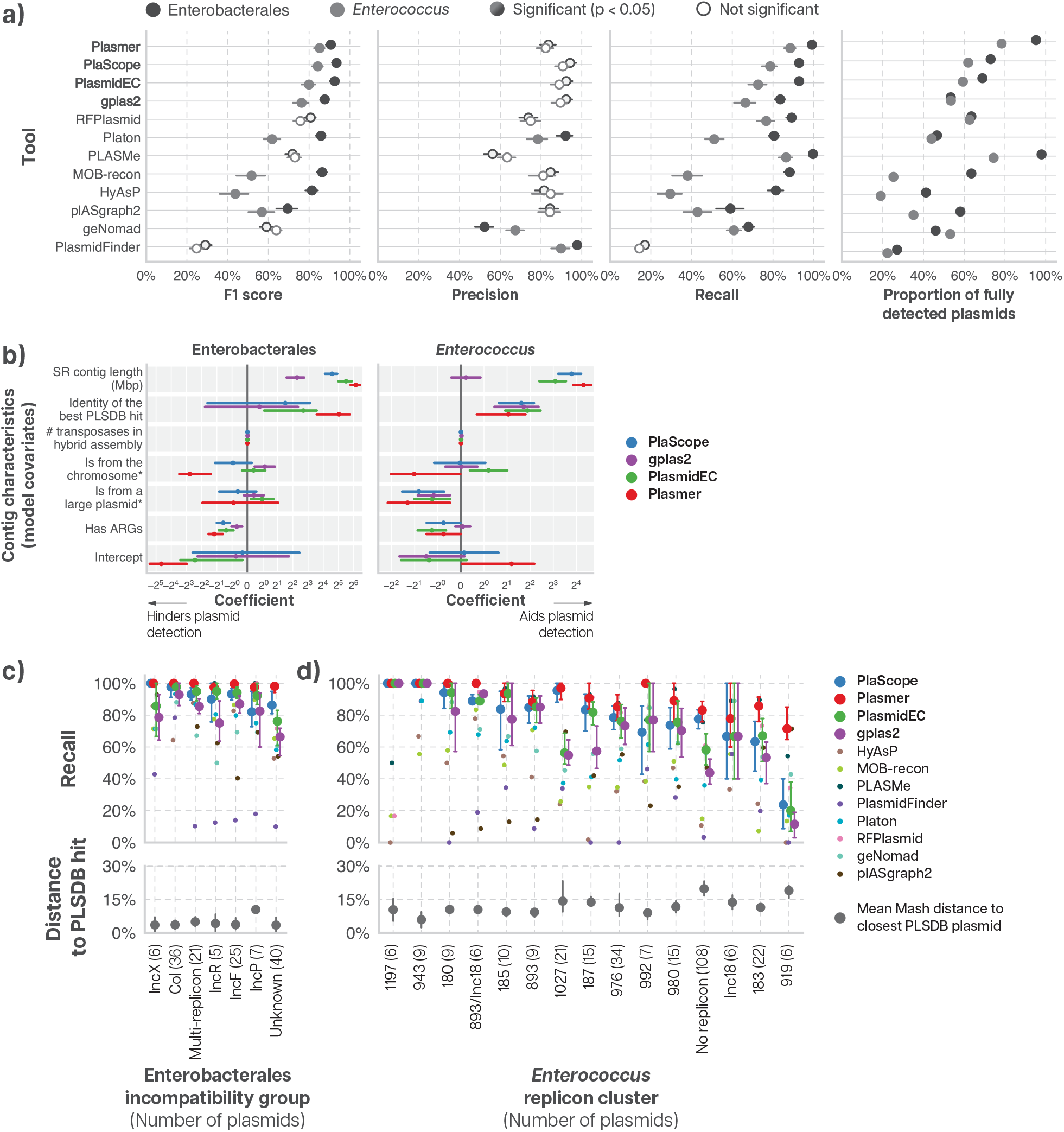
Representation in plasmid databases is important for accurate plasmid detection, making enterococcal plasmids harder to detect. **a)** Plasmid detection metrics for Enterobacterales and *Enterococcus*: F1 score, precision, recall, and the proportion of fully detected plasmids (i.e., all short-read contigs correctly classified as plasmid). The lines show the 95% confidence interval, and solid points represent significant differences between taxa (two-sided bootstrapping test with Benjamini-Hochberg correction). **b)** We trained logistic regression models to assess the individual effects of contig characteristics on plasmid detection. This figure shows the coefficients for specific assembly and plasmid characteristics for the four best plasmid detection tools. Characteristics with positive coefficients aid plasmid detection, while those with negative coefficients hinder it. Lines, colored by tool, represent 95% confidence intervals. ^*^ Relative to small plasmids (contigs were grouped into three classes: chromosomal, small plasmid, and large plasmid). **c-d)** Plasmid detection recall (top panel) for **c)** Enterobacterales incompatibility groups and **d)** *Enterococcus* replicon clusters, for groupings with at least five plasmids. We grouped plasmids with finished hybrid assemblies according to their replicon cluster, and further grouped Enterobacterales replicon clusters into incompatibility types to aid interpretation. This further grouping could not be done for enterococcal plasmids, as most had no associated incompatibility type. The bottom plots show the mean Mash distance to the most similar plasmid in PLSDB for each incompatibility group/replicon cluster. Error bars show the 95% confidence intervals for the best four tools.

##### Contiguity and representation in databases are important for plasmid detection

We hypothesized that plasmid detection would be affected by plasmid and assembly characteristics, such as plasmid size, fragmentation, or the presence of mobile elements. Since the individual effects of these characteristics are difficult to disentangle, especially as some are correlated (e.g., ARG-containing plasmids are often larger and contain more mobile genetic elements [38, 39]), we used logistic regression models to determine the individual influence of five assembly and plasmid characteristics on the performance of each of the four best plasmid detection tools (Methods; Figure 3b). We then explored the characteristics that significantly affected plasmid detection in greater detail by comparing results for plasmids with these characteristics against the results for those without.

###### Plasmids represented in databases were easier to detect

Enterobacterales plasmids were easier to detect than the mostly novel *Enterococcus* plasmids (highest F1 score of 93.5% versus 84.2%, respectively), mainly due to a greater fraction of Enterobacterales plasmid contigs being correctly identified as plasmidic (Figure 3a, Supplementary Table 5). This was despite the fact that the enterococci assemblies were less fragmented than their Enterobacterales counterparts. For instance, Plasmer, the best tool for the enterococci, failed to detect 11.4% of *Enterococcus* versus 0.9% of Enterobacterales plasmid contigs, corresponding to 5.5% and 1.0% of plasmid base pairs, respectively. Furthermore, Plasmer correctly classified all contigs from 95.3% of all Enterobacterales plasmids, but only 78.2% of *Enterococcus* plasmids (Figure 3a, Supplementary Table 6). This difference was mainly driven by database representation, as indicated by our logistic regression model (Figure 3b): Enterobacterales plasmids were better characterized in plasmid databases and thus easier to detect.

This pattern was also observed at the level of incompatibility or replicon types (Methods; Figure 3c-d): plasmids from replicon types that were more difficult to detect were also more dissimilar to plasmids in PLSDB (Figure 3d, Supplementary Table 7-9). In particular, all four tools had difficulty detecting *Enterococcus* plasmids from rep cluster 919, possibly because these did not match plasmids in the PLSDB database well nor any sequence in the NCBI Nucleotide database with >25% coverage.

###### Large plasmid contigs from less fragmented assemblies were easier to detect

As expected, the logistic regression model suggested that assembly contiguity improved plasmid detection, with longer contigs being easier to detect (Figure 3b, Supplementary Table 10). Larger individual contigs also led to improved performance across plasmid detection tools, particularly for the enterococci, primarily due to increased recall (Figure 4a; Supplementary Figure 2). Somewhat counterintuitively, small (*<* 28 kbp) *Enterococcus* plasmids (Supplementary Figure 3) were easier to detect (higher recall) than larger ones (Figure 4b); however, these smaller plasmids were also highly contiguous in our short-read assemblies: 93.3% of small enterococcal plasmids were complete and in a single contig, whereas only 37.8% of large plasmids were complete (Supplementary Table 11). Enterobacterales plasmid recall was similar across short and long plasmids (Figure 4b).

**Figure 4.**
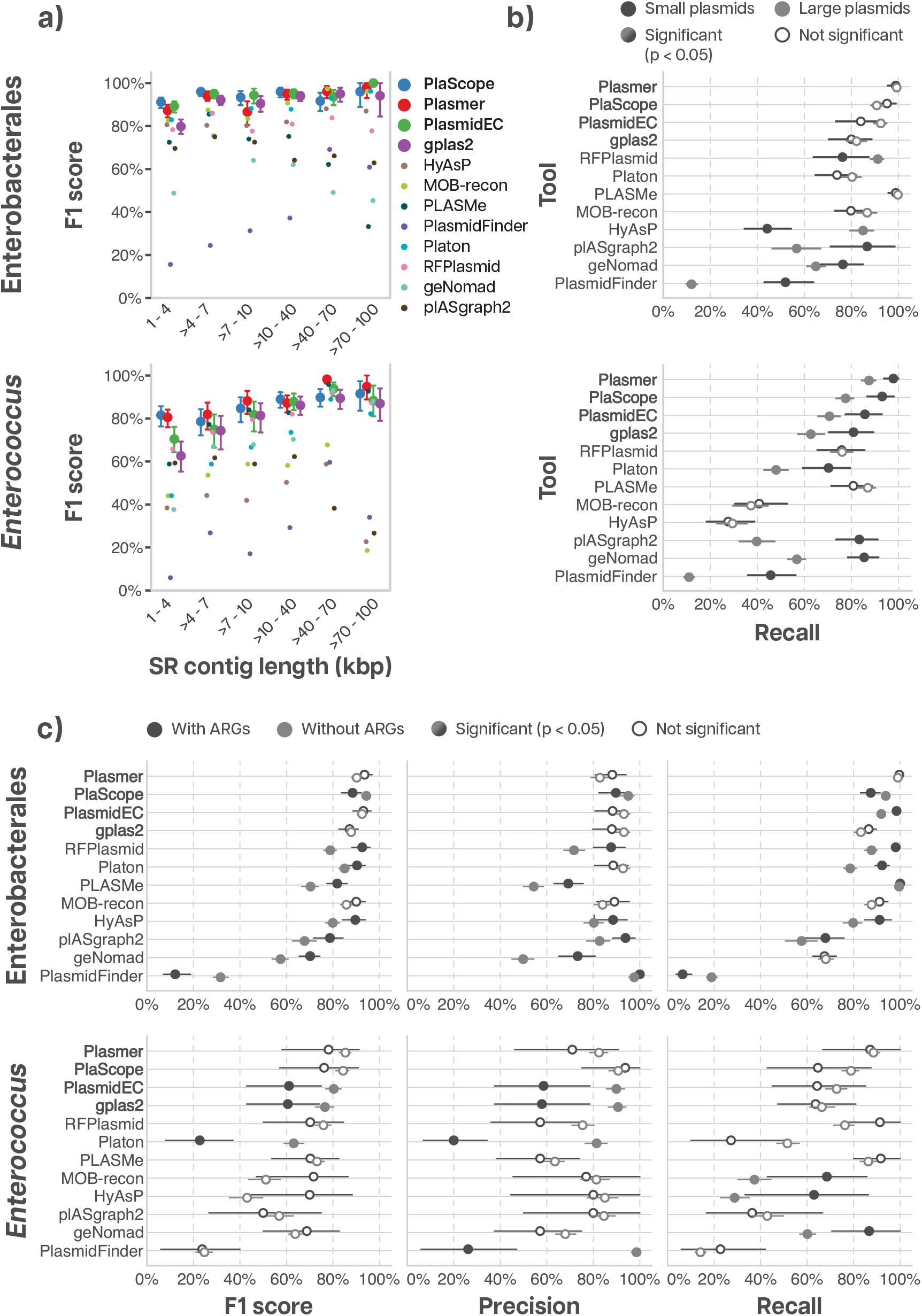
SR contig length, plasmid size, and presence of ARGs affect plasmid classification. **a)** F1 scores (colored markers) as a function of SR contig length. **b)** Recall for contigs from small vs. large plasmids. **c)** Plasmid detection metrics for SR contigs with or without ARGs. In all panels, error bars represent the 95% confidence interval. In b) and c), full points denote significant differences between groups (two-sided bootstrapping test with Benjamini-Hochberg correction).

###### Some tools found plasmids with antimicrobial resistance genes more difficult to detect

In our dataset, Enterobacterales plasmids were more likely to carry ARGs (39% of plasmids, with an average of 2.4 ARGs per plasmid) than those from the enterococci (6%, average of 0.1 per plasmid), possibly due to their differing origins (clinical vs. non-human-associated, respectively) (Supplementary Tables 12-13). Plasmids with ARGs had more transposases, which can induce breaks in the assemblies, leading to shorter contigs that are more difficult to classify [38, 39] (Supplementary Figure 4). Plasmids without ARGs were easier to detect, though this trend was not consistently observed across all tools (Figure 4c, Supplementary Table 14).

#### Plasmid reconstruction

##### Gplas2 was the best plasmid reconstruction tool, but few plasmids were perfectly reconstructed

Since few short-read assembled plasmids are represented by a single contig, it was also of interest to understand how well tools for plasmid reconstruction (or binning) perform in assigning different contigs to the same plasmid. We ran four plasmid reconstruction tools (Table 1) end-to-end, using their own built-in plasmid detection utility, followed by plasmid reconstruction. We quantified the quality of predictions according to their i) homogeneity, which reflects the purity of the predicted plasmid bins; ii) completeness, which reflects whether bins include all of the short-read contigs from a plasmid; and iii) *Normalized Mutual Information* (NMI), which combines homogeneity and completeness (Methods). We only compared three of the four tools for our *Enterococcus* dataset, because PlasBinFlow (using default settings) failed to detect any plasmid contigs in enterococci.

Gplas2 outperformed the other tools for both Enterobacterales and the enterococci (Figure 5a, Supplementary Table 15); however, despite the high NMI, homogeneity, and completeness, less than half of the plasmids - 47.3% (n=70) and 49.2% (n=151) of Enterobacterales and *Enterococcus* plasmids, respectively - were perfectly reconstructed (i.e., all contigs from the same plasmid binned together without contamination from contigs from other plasmids or the chromosome). Further, complete plasmids made up of just one 5 contig were frequently misassigned to another plasmid bin, with 23.1% (n=16) of complete Enterobacterales and 19.7% (n=31) of complete *Enterococcus* plasmids binned together with contigs from other plasmids. When excluding complete plasmids, gplas2 perfectly reconstructed only 21.5% (n=17) of Enterobacterales and 16.8% (n=25) of *Enterococcus* plasmids.

**Figure 5.**
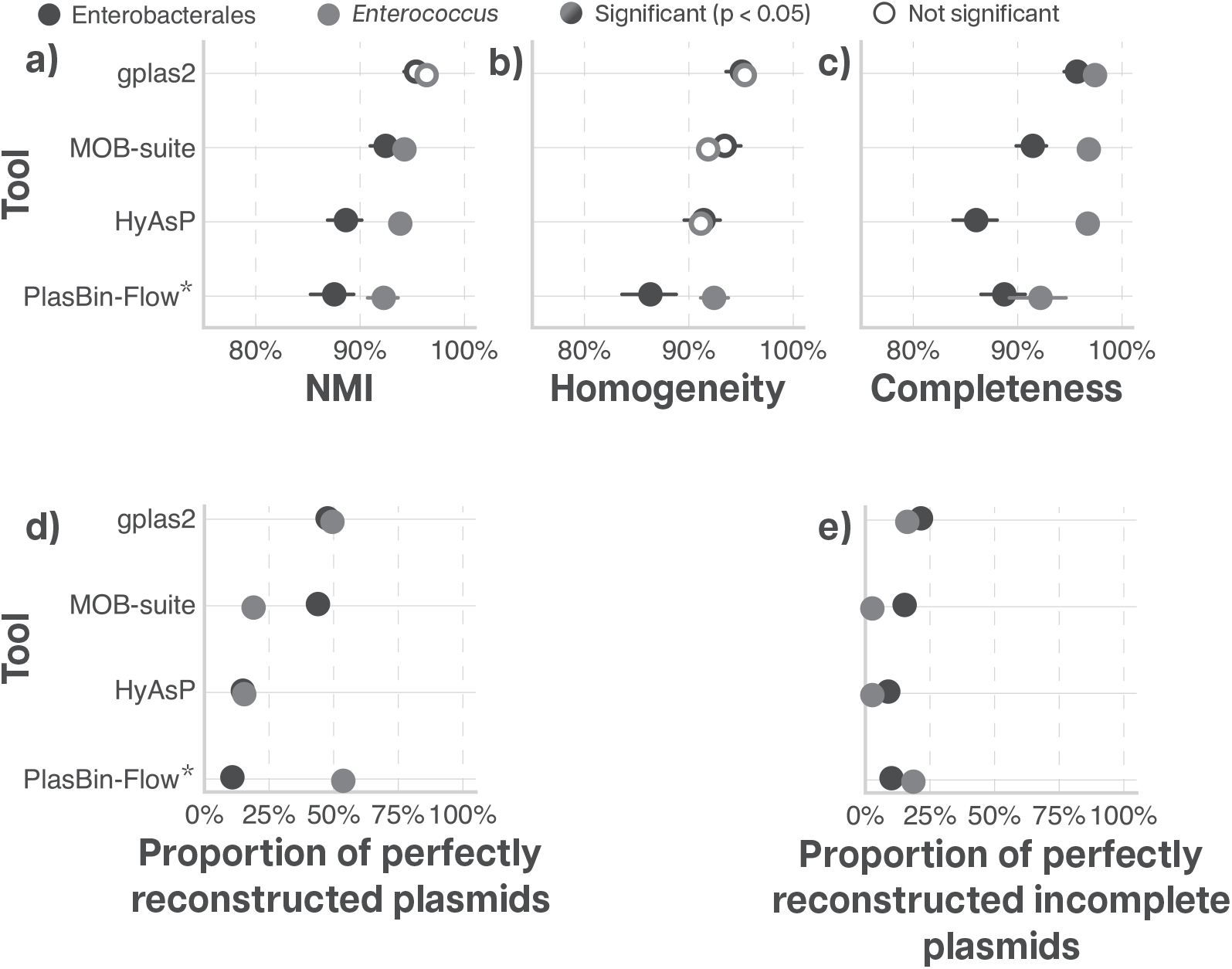
Few plasmids were perfectly reconstructed. Plasmid reconstruction metrics for Enterobacterales and enterococci, which had more contiguous assemblies: Normalized Mutual Information (NMI), homogeneity, completeness, the proportion of perfectly reconstructed plasmids (i.e., all contigs from a plasmid binned together with no contamination from other plasmids), and the proportion of perfectly reconstructed incomplete plasmids (i.e., fragmented into >1 contig in the short-read assembly). ^*^ To reconstruct enterococcal plasmids with PlasBin-Flow, we used the contig annotations from Plasmer, the best plasmid detection tool for enterococci. Error bars show the 95% confidence interval, while full points represent significant differences between taxa (two-sided bootstrapping test with Benjamini-Hochberg correction).

To control for variability in the plasmid detection component of each tool, we also reran PlasBinFlow and gplas2 using the contig classifications from the best plasmid detection tools (PlaScope for Enterobacterales and Plasmer for the enterococci) as input for the plasmid reconstruction step. HyAsP and MOB-suite could not perform the detection and reconstruction steps independently. For gplas2, this approach did not lead to any significant improvements over running it end-to-end; however, the reconstruction of *Enterococcus* plasmids with PlasBin-Flow improved significantly, with an NMI similar to that of MOB-suite and HyAsP end-to-end, but still worse than gplas2 (Supplementary Figure 5 and Supplementary Table 16).

##### Higher contiguity improves plasmid reconstruction

We next used a linear regression model to assess the individual effects of specific assembly characteristics on plasmid reconstruction by modeling their impact on NMI (Methods). Assembly fragmentation had a significant effect, with more fragmented assemblies (p *<* 0.001, Supplementary Table 17) and those with more plasmids (p = 0.001) leading to greater difficulty in reconstructing plasmids. Surprisingly, transposases were associated with better plasmid reconstruction, albeit with a small effect size. However, only a small proportion of the variation in NMI was explained by the assembly characteristics included in this analysis (R^2^ = 0.177 for the training set), so there are likely additional features that impact plasmid reconstruction.

Overall, NMI was slightly greater for the enterococci than for Enterobacterales; however, this difference was not significant for gplas2 (Supplementary Table 18). Although we found no differences in homogeneity between taxa, the reconstructed *Enterococcus* plasmids had greater completeness. This can be attributed to the higher contiguity of *Enterococcus* plasmids in short-read assemblies when compared to Enterobacterales, making it easier to bin most of a plasmid together.

Though ARGs did not significantly affect plasmid reconstruction based on our linear regression models, we found that ARG-containing plasmids, which are more fragmented (Supplementary Figure 4), were more difficult to reconstruct when comparing reconstruction metrics separately for plasmids with and without ARGs (Figure 6). Enterobacterales plasmids with ARGs were more difficult to reconstruct for three of the four tools tested, including gplas2, the best plasmid reconstruction tool (Figure 6, Supplementary Table 19). We saw a similar trend for the enterococci, although this difference was not significant.

**Figure 6.**
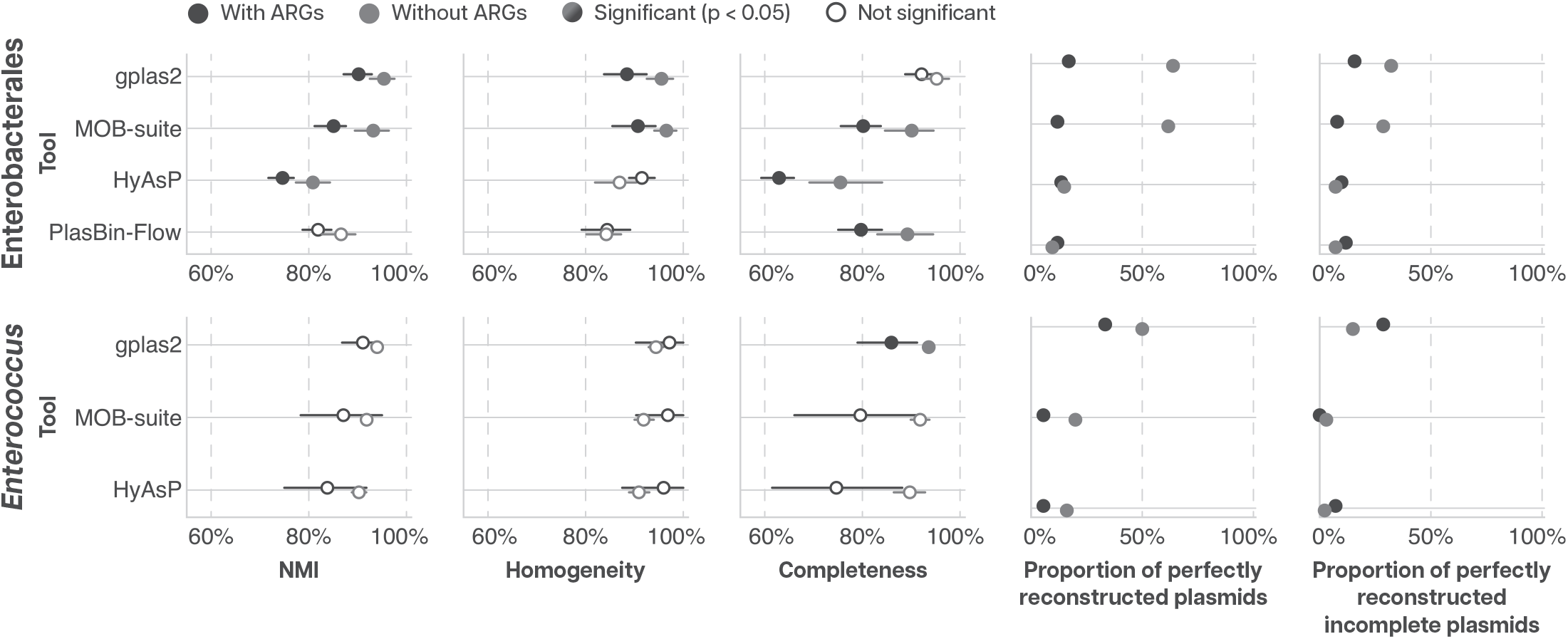
Some tools had more difficulties reconstructing plasmids with ARGs. Plasmid reconstruction metrics for plasmids with and without ARGs. We grouped plasmids according to ARG content and calculated the reconstruction metrics for each group. Error bars show the 95% confidence interval, while full points represent significant differences between ARG-containing and non-ARG-containing plasmids (two-sided bootstrapping test with Benjamini-Hochberg correction).

## Discussion

In this study, we compared tools for plasmid detection and reconstruction from short-read assemblies. We established a challenging benchmarking dataset, examining a broader diversity of genomes and plasmids than used in previous studies benchmarking plasmid detection and reconstruction tools, which primarily focused on *E. coli* [40, 11, 12].

We included 13 state-of-the-art tools, which use different methodologies. Plasmid detection was very good overall (the best tools for Enterobacterales and the enterococci detected 93% and 89% of plasmid contigs, respectively, with just 6% and 18% of false positives), but four tools stood out as the best: Plasmer, PlaScope, PlasmidEC, and gplas2. These use a variety of data types (contigs or assembly graphs) and methodologies (purely reference-based or with machine learning), demonstrating that different computational strategies can yield high performance.

Plasmid detection tools achieved better results for plasmids that were better represented in plasmid databases, which are used by many of the tools: Enterobacterales plasmids were easier to detect than those from the enterococci, despite being more fragmented, and the tools had more difficulty detecting plasmids of less well characterized replicon types. The more similar a plasmid was to known plasmid sequences in the PLSDB database, the more likely it was to be detected, after adjusting for other characteristics of the short-read assemblies. Most of the enterococci included in our dataset were collected from feces of undomesticated animals, including insects in remote locations, and harbored many highly diverse plasmids with little representation in plasmid databases, which are biased towards plasmids from human-associated bacteria, largely of clinical origin [16]. We did not include plasmids from clinically-adapted enterococci so as to assess how the tools behaved for novel plasmids. As a consequence, most plasmids from our set of enterococci had no matches to sequences in PLSDB (although the mosaic structure of plasmids makes it difficult to define if a plasmid is present in a database or not). Plasmids from human-adapted *E. faecalis* and *E. faecium* would likely be detected more easily, as commonly occurring plasmids from hospital-adapted enterococcal strains are well-represented in PLSDB. As long-read sequencing becomes more accessible, expanding plasmid databases with novel and high-quality sequences may improve the future performance of current plasmid detection tools.

As expected, longer contigs and less fragmented assemblies led to better plasmid detection results, possibly because longer contigs are more likely to contain typical plasmid features, and have at least partial matches to sequences in databases. This trend was more pronounced for our enterococci, maybe because they were less well characterized in plasmid databases and required longer sequences to yield matches to known plasmid content. We also found that small *Enterococcus* plasmids were more easily detected than large ones. Our small *Enterococcus* plasmid assemblies were unusually contiguous, which aided in their identification, and it is also possible that the small plasmids in our diverse dataset are less variable and better described than long ones, making them easier to detect [41]. Importantly, the *Enterococcus* short-read assemblies were of very high contiguity relative to those from Enterobacterales, which may not be representative of all short-read assemblies. Given how impactful we found fragmentation to be on plasmid detection, the tools included may not perform as well for *Enterococcus* assemblies with lower quality.

Plasmid reconstruction, or binning contigs identified as plasmidic into discrete plasmids, was very challenging for existing tools. This highlights the need for improvements in methods for reconstructing plasmids using short-read datasets. Gplas2 was the best plasmid reconstruction tool for both taxa, despite perfectly reconstructing less than half of the plasmids. Gplas2 uses a similar approach to PlasBin-Flow and HyAsP, but it allows for a greater variance in the read coverage for each predicted plasmid, which may improve results. PlasBin-Flow failed to detect any *Enterococcus* plasmids prior to the plasmid reconstruction step, as its default database contained plasmids from very few enterococcal isolates [22]. However, PlasBin-Flow includes the functionality to create custom reference databases, which would likely improve its performance with enterococcal plasmids.

Using Gplas2, plasmids reconstructed from the enterococci had higher completeness than those from Enterobacterales, reflecting the higher contiguity of our *Enterococcus* short-read assemblies. Unsurprisingly, plasmids in more fragmented assemblies were more difficult to reconstruct, a result that held even when adjusting for the number of plasmids and chromosomes in the assembly, and the number of ARGs and transposases. Unlike plasmid detection, the performance of plasmid reconstruction tools was similar between taxa based on NMI. This is likely because plasmid reconstruction is not as dependent on the breadth of diversity captured by sequence databases, but more on the topology of the assembly graph [42]. To ensure plasmids are reconstructed accurately, sequencing and assembly methods should be chosen carefully to maximize contiguity.

As studying the spread of antimicrobial resistance is one of the main motivators for studying plasmids, it is critical to understand the impact of ARGs on the ability to classify contigs as plasmidic and bin contigs together into plasmids. A previous comparison found that large plasmids with ARGs were as easy to detect as those without [12]. However, this study used a different definition of detection than we did: plasmids were only considered undetected if none of their contigs were annotated as plasmidic, while we assessed plasmid detection on a contig-by-contig basis.

Globally, we saw the same trend of ARGs not affecting plasmid detection in our dataset when separately analyzing plasmids with and without ARGs, but some tools had more difficulty in classifying contigs with ARGs. Our logistic regression models also identified contigs with ARGs to be more difficult to classify. ARGs are often within small mobile elements that induce breaks in the assemblies, and thus are located within shorter contigs [38, 39]. Because shorter contigs are more difficult to classify and these small mobile elements can also be integrated in the chromosome, some tools may misclassify contigs with ARGs more often. While transposases had a negligible effect on plasmid detection according to our logistic regression models, these models may have been unable to accurately disentangle the individual effects of ARGs and transposases given their strong association. Using several tools together may aid in detecting plasmids, particularly when looking for ARG-containing plasmids; however, PlasmidEC, an ensemble of tools, also had more difficulty in detecting ARG-containing plasmids from the enterococci.

In terms of plasmid reconstruction, Enterobacterales plasmids with ARGs were more difficult to reconstruct for most tools, including the highest-scoring tool, gplas2, which was also observed in previous comparisons [12]. ARGs are associated with transposable elements with multiple copies in the genome which may induce breaks in the assembly [38], making plasmids with ARGs more difficult to reconstruct. Indeed, plasmids with ARGs were larger and more fragmented in our short-read assemblies. Although our linear regression model found transposases to slightly aid plasmid reconstruction, the effect size was negligible, and there may be other factors influencing plasmid reconstruction that were not included in the model.

Among the limitations of our work, although we tested tools with a greater diversity of isolates than previous comparisons, including plasmids from highly diverse, non-clinical enterococci from remote locations, our benchmarking dataset still only covered a small fraction of all bacterial diversity. Most trends observed in this work were not specific to a taxon; nevertheless, plasmid detection and reconstruction may be more challenging for taxa less well-studied than Enterobacterales or *Enterococcus*, which would have even less representation in plasmid databases.

Also, we focused on the most recent tools in this benchmarking work, as well as older highly-cited tools. It is possible that some other older, well-cited tools that we did not include may be suitable alternatives in some settings [43, 44, 45, 46], but these have previously been benchmarked and were found to have a poorer performance than tools we included in this study [47, 12].

Finally, data from long-read sequencing technologies has become the gold standard for characterizing plasmid sequences, though these data sometimes struggle to produce small plasmids [10, 48]. Long-read sequencing will likely continue to become more accessible and increasingly widely used when analyzing plasmids. But detecting and reconstructing plasmids from short-read data remains important: generating short-read data is still considerably cheaper, and the vast majority of the assemblies in public databases are generated from short reads.

## Conclusions

In our comparison of methods to detect plasmids based on short-read assemblies using a diverse set of bacterial species, we found that PlaScope, Plasmer, PlasmidEC, and gplas2 stood out as the best tools. However, it was more difficult to detect plasmids from species that are not well represented in plasmid databases. For less well-characterized species, augmenting databases with additional plasmid information tailored to these species would help identify plasmids. In our comparison of methods to reconstruct plasmids, we found that this task was challenging, even for the highest-performing tool (gplas2). We found that assembly contiguity was crucial for plasmid reconstruction, which made plasmids with ARGs (which are usually more fragmented) more difficult to reconstruct. This benchmarking study will help researchers choose the best tool to detect and reconstruct plasmids from short-read assemblies in future studies, and highlights key areas for future improvement for plasmid detection and reconstruction tools.

## Data availability

The data underlying this article are available in the Sequence Read Archive at https://www.ncbi.nlm.nih.gov/sra, the BioSample database at https://www.ncbi.nlm.nih.gov/biosample (and can be accessed with the accession numbers in Supplementary Table 1), and in the article and in its online supplementary material. The code used for analysis is available at https://github.com/broadinstitute/plasmid-detection-benchmark.

## Acknowledgments

This work was supported by Federal funds from the National Institute of Allergy and Infectious Diseases, National Institutes of Health, Department of Health and Human Services, under Grant Numbers U19AI110818 to the Broad Institute and U19AI172725 to University of California Irvine and the Broad Institute.

## Notes

### Competing Interest Statement

The authors have declared no competing interest.

### Summary of Updates

The author list was updated, Terrence Shea has been added

