## Supplementary Figure for "Circling in on plasmids: benchmarking plasmid detection and reconstruction tools for short-read data from diverse species"

### Supplementary Figures

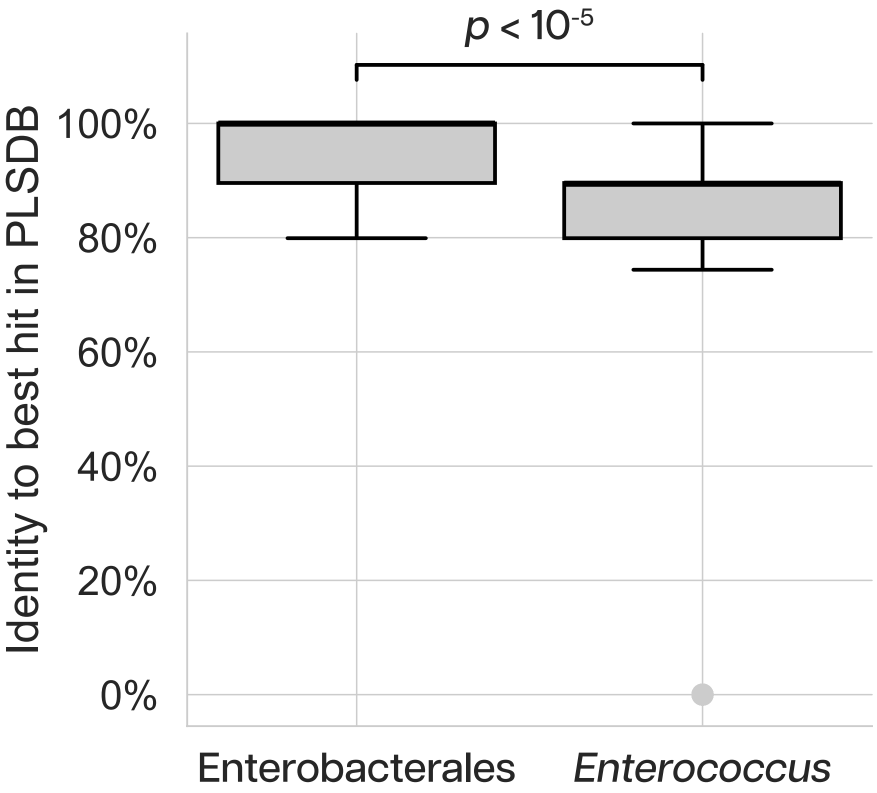


Supplementary Figure 1: **Enterobacterales plasmids were better represented in PLSDB than those from enterococci.** Distributions of the identity between each plasmid with a finished hybrid assembly and its best hit in PLSDB, calculated using Mash.

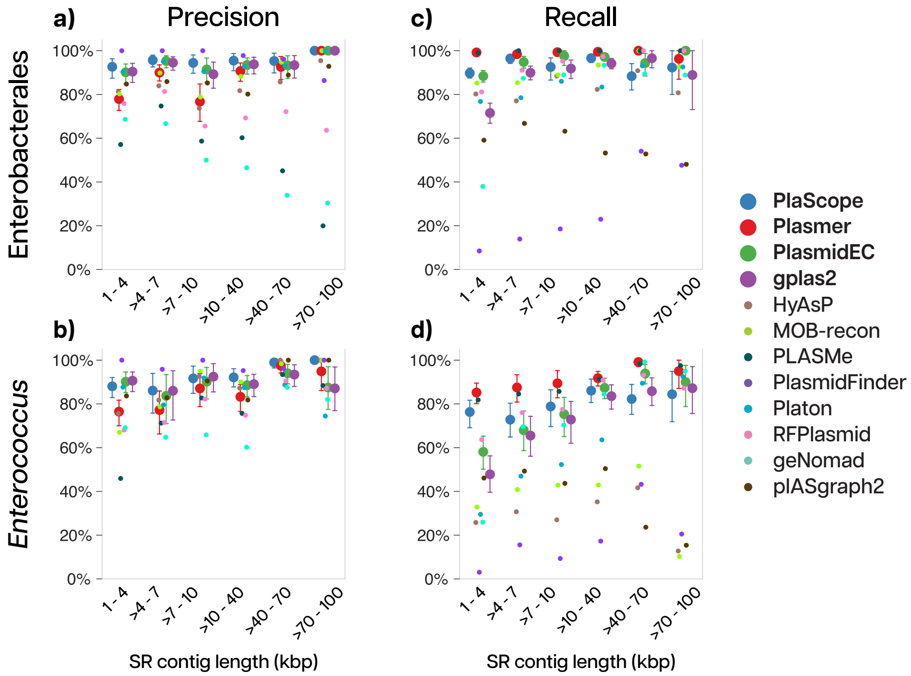


Supplementary Figure 2: **Longer plasmid contigs were easier to detect.** **a-b)** Precision and **c-d)** recall for plasmid detection as a function of short-read contig length; for **a, c)** Enterobacterales and **b, d)** *Enterococcus*. Error bars show the 95% confidence interval for the four best tools.

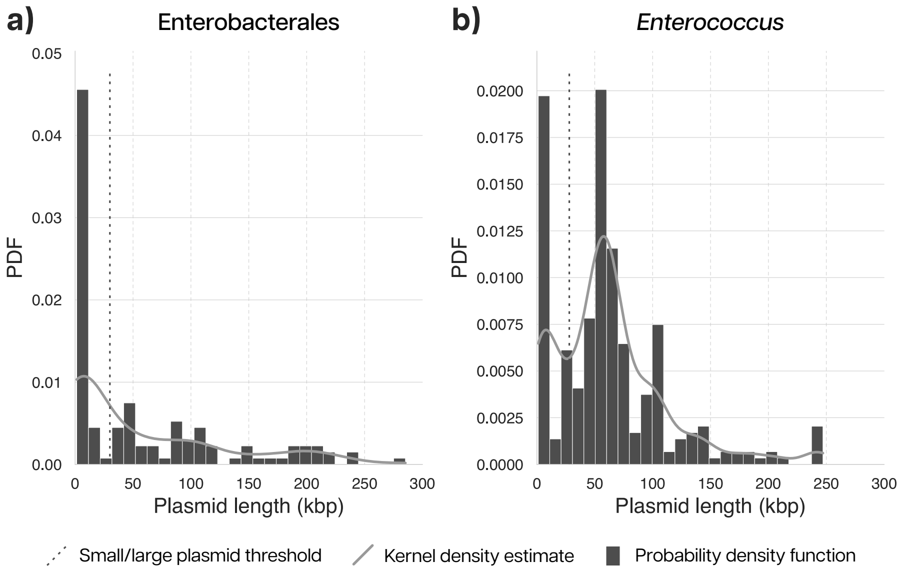


Supplementary Figure 3: **Plasmids in our dataset have a bimodal length distribution.** Histogram (bars) and kernel density estimate (solid line) of plasmid lengths for complete plasmids in the hybrid assemblies from **a)** Enterobacterales and **b)** enterococci. The dashed lines show the threshold used to classify plasmids as “small” or “large”.

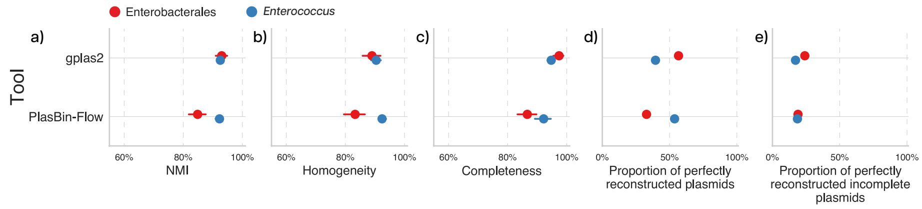


Supplementary Figure 4: **Using optimized plasmid classifications as input to plasmid reconstruction improves the performance of PlasBin-Flow significantly**. Plasmid reconstruction metrics for Enterobacterales and *Enterococcus* using the plasmid detection results from the best detection tools (PlaScope for Enterobacterales and Plasmer for *Enterococcus*): **a)** Normalized Mutual Information (NMI), **b)** homogeneity, **c)** completeness, **d)** the proportion of perfectly reconstructed plasmids (i.e., all contigs from a plasmid binned together with no contamination from other plasmids), and **e)** the proportion of perfectly reconstructed incomplete plasmids (i.e., fragmented into >1 contig in the short-read assembly). Error bars represent the 95% confidence interval.

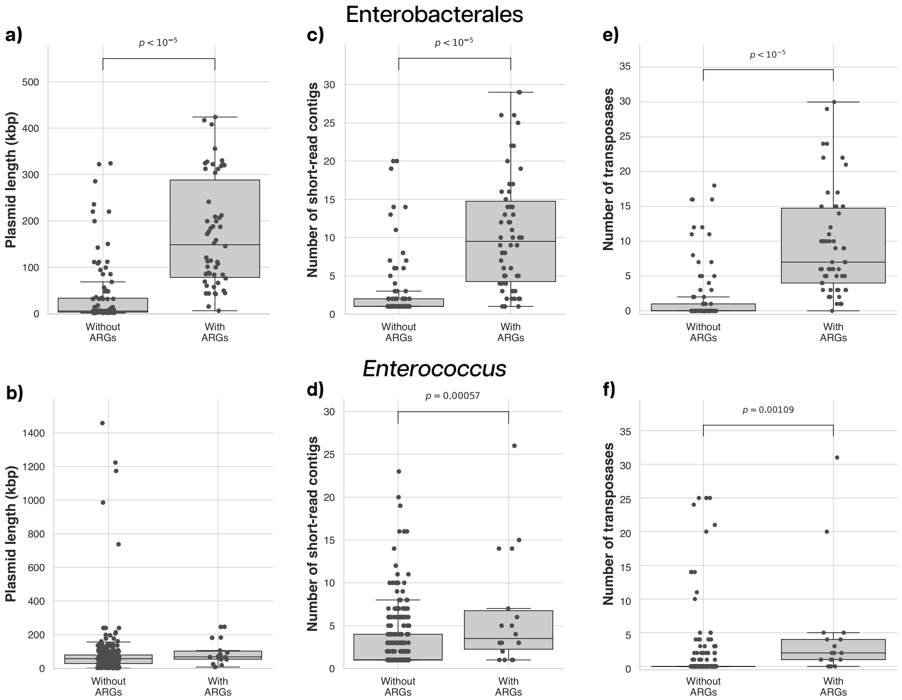


Supplementary Figure 5: **Plasmids with ARGs were larger and more fragmented than those without ARGs**. **a-b)** Distribution of lengths and **c-d)** the number of short-read contigs for **a, c)** Enterobacterales and **b, d)** enterococcal plasmids with and without ARGs. **e-f)** Distribution of the number of transposases in plasmids with and without ARGs for **e)** Enterobacterales and **f)** *Enterococcus*. Bars denote significance (two-sided t-test for independent samples).
